# Single-cell analysis identifies TOP2α as a critical regulator of G2/M entry and differentiation-dependent productive HPV replication

**DOI:** 10.64898/2026.09.23.753843

**Authors:** Arushi Vats, Olga Rozhok, Laimonis Laimins

## Abstract

High-risk human papillomaviruses (HPVs) are the etiological agents of over 5% of cancers worldwide including those of the cervix and oropharynx. HPVs infect stratified epithelia and establish latent infections in basal cells but restrict productive replication or amplification to differentiated suprabasal cells. Despite the presence of viral genomes in most differentiated cells, only a subset of cells amplifies genomes as well as express late genes and these correspond to those that have re-entered G2/M, but the factors regulating this selectivity are unknown. To determine the signals that control the productive viral life cycle, single cell RNA seq was performed on cells that stably maintain high-risk HPV genomes following differentiation. Ten populations of undifferentiated and differentiating keratinocytes were identified, however, only one differentiated population had entered G2/M, expressed late genes and amplified viral genomes. One of the highly expressed replication factors in this population was the type II topoisomerase TOP2α while no other topoisomerases were similarly induced. TOP2α was found to bind to viral genomes, and acute depletion in differentiating cells blocked entry into G2/M, impaired genome amplification and abrogated late gene expression. Amplification also requires activation of DNA repair pathways through the induction of high levels of DNA breaks, and TOP2α accounted for more than half of the breaks present in differentiating cells. Increases in levels of TOP2α in differentiating cells were driven by the E7 oncoprotein acting through the transcription factor FOXM1, which also controls expression of G2/M factors suggesting an auto-regulatory loop. These studies identify TOP2α as a critical regulator of HPV genome amplification upon differentiation.

## Introduction

Human papillomaviruses (HPVs) are the causative agents of cervical cancer and of most oropharyngeal cancers^1,2^. High-risk HPVs infect cells in the basal layer of stratified epithelia and establish their genomes there as low-copy nuclear episomes. Productive replication, or amplification, is restricted to a discrete subset of differentiated cells in the suprabasal layers and the signals regulating this restriction are not known. The majority of sexually active individuals acquire a genital HPV infection during their lifetime^3^, and most clear it within a few years^4^. In the subset of individuals that fail to clear, persistent lesions are established that can progress to cancer, a transition frequently accompanied by integration of viral sequences into host DNA and loss of the capacity to produce progeny virions^5^. Defining the signals that govern the productive phase of the life cycle can identify the factors that permit progression. Viral genomes are replicated in undifferentiated cells during S phase, in step with cellular replication, whereas amplification takes place in suprabasal cells that have re-entered G2/M^6–8^. Entry into G2/M is required because HPV amplification depends on activation of the ATM and ATR DNA damage pathways, which require a homologous DNA template for repair^9–13^. While viral genomes are present in all differentiating suprabasal cells only a subset re-enter G2/M for amplification, and the present study examines the factors responsible for this restriction.

In normal stratified epithelia, cells arrest in G0 as they leave the basal layer to undergo differentiation. In contrast, HPV-positive cells arrest in G1 as they begin to differentiate, and a subset pass through S phase to enter G2/M in the suprabasal layers to allow for amplification^7,8^. This cell cycle progression is driven by the viral oncoproteins E6 and E7. E7 targets the retinoblastoma protein Rb for degradation, releasing E2F factors from repression, while E6 binds p53 and directs its rapid turnover^14–17^. E6 and E7 are also responsible for activating the DNA repair pathways that are required for amplification^9,10,13^. The late phase of the viral life cycle includes high-level expression of the fusion protein E1^E4, which is synthesized specifically in cells undergoing amplification. E1^E4 binds to cytokeratin networks and induces their collapse to allow newly assembled virions to escape^18,19^. E1^E4 is the most abundantly expressed viral protein and is found specifically in cells undergoing amplification^20^.

The differentiation-dependent life cycle can be reproduced in culture using cells derived from cervical biopsies or generated by transfection of human keratinocytes with recircularized genomes that can be induced to differentiate either by a switch in calcium concentration or following growth in organotypic cultures^21–25^. To identify the signals that specifically regulate re-entry into G2/M and to induce amplification in differentiating cells, single-cell RNA sequencing of differentiated HPV31-positive cells that stably amplify episomes was performed. Ten clusters of undifferentiated and differentiated HPV positive cells were identified, but only one arose within the differentiated compartment, had re-entered G2/M and expressed high levels of E1^E4 together with other viral genes. These cells were also shown to amplify HPV genomes. Among the most prominent replication factors increased specifically in the G2/M cluster was the type II topoisomerase TOP2α. Topoisomerases modulate higher-order chromatin structure by transiently breaking and religating one or both strands of duplex DNA^26,27^. Type I enzymes (TOP1, TOP3α and TOP3β) cleave a single strand, whereas type II enzymes (TOP2α and TOP2β) cleave both. Levels of both type I and type II topoisomerases are elevated in undifferentiated HPV-positive cells relative to uninfected keratinocytes^28,29^, but only TOP2α was induced in the differentiating population that re-entered G2/M and was undergoing amplification. Depletion of TOP2α specifically in differentiating cells suppressed the appearance of cells in G2/M and blocked expression of factors such as CDK1, which prevented HPV genome amplification. Induction of TOP2α was controlled by E7 acting through FOXM1, the transcription factor that helps drive the expression of genes in the G2/M cluster^30–32^. These studies identify TOP2α as a critical regulator of HPV genome amplification upon differentiation.

## Results

### Single-cell profiling identifies an HPV amplification-competent state

In order to investigate the signals regulating the productive phase of the HPV lifecycle, calcium-induced differentiation of HPV positive keratinocytes was used as it faithfully recapitulates amplification and late gene expression^21,25^. HPV31-positive CIN612 9E keratinocytes were derived from a low-grade cervical biopsy and stably maintain approximately 50 copies of viral episomes^22,23^. These cells were differentiated by sequential calcium switch and harvested after 96 h (144 h total; **Fig. 1a**). This endpoint was selected because induction of the spliced *E1^E4* late viral transcript, which marks entry into the productive program^19,20^, was increased 11.5-fold over undifferentiated cells, keratin 10 was strongly induced and cell viability remained at 90% (**Supplementary Fig. 1**). Single cell RNA-seq was performed on undifferentiated (UD) and differentiated (D4) cultures and analyzed in parallel on the 10x Genomics Chromium platform. Since viral transcripts constitute a small fraction of reads in this system, reads were aligned to a combined GRCh38–HPV31 reference with viral open reading frames annotated as individual features and reads assigned to low-confidence barcodes were retained; 24,059 cells passed quality control.

**Figure 1.**
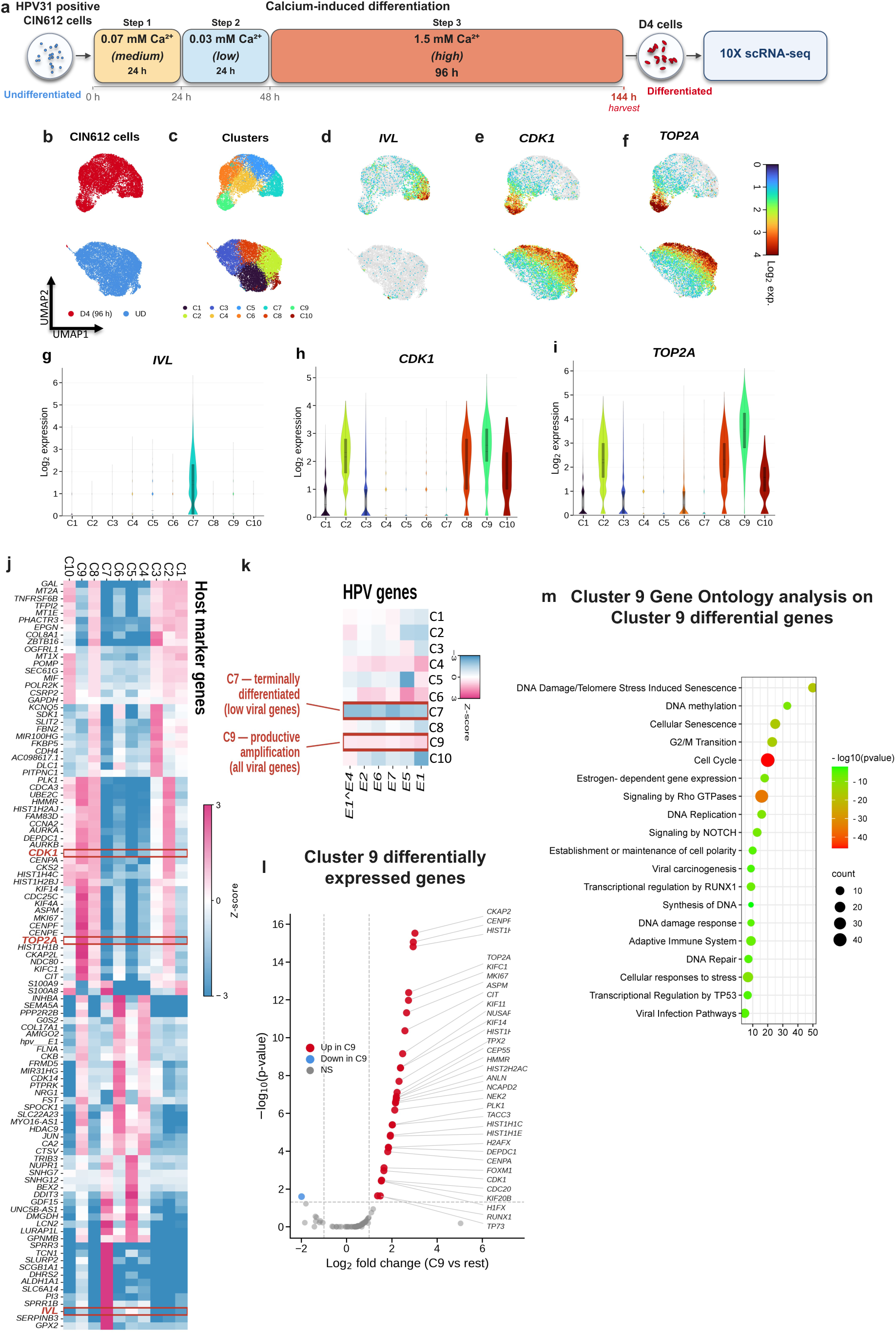
Single-cell RNA sequencing of differentiating HPV31-positive keratinocytes identifies an amplification-competent G2/M population. a, Experimental scheme. HPV31-positive CIN612 9E keratinocytes were differentiated by stepwise calcium addition (0.07 mM, 24 h; 0.03 mM, 24 h; 1.5 mM, 96 h) and harvested at 144 h, and undifferentiated (UD) and differentiated (D4) cultures were processed in parallel for 10x Genomics single-cell RNA sequencing. **b**, UMAP embedding of 24,059 cells coloured by condition (D4, red; UD, blue). **c**, The same embedding coloured by the ten transcriptional clusters (C1–C10). **d–f**, Feature plots showing log₂ normalized expression of *IVL* (**d**), *CDK1* (**e**) and *TOP2A* (**f**); D4 cells upper, UD cells lower. **g–i**, Violin plots of *IVL* (**g**), *CDK1* (**h**) and *TOP2A* (**i**) expression across clusters C1–C10. **j**, Heat map of marker gene expression (row-scaled *z*-score) across clusters; *CDK1*, *TOP2A* and *IVL* are boxed. **k**, Heat map of HPV31 viral gene expression (*E1^E4*, *E2*, *E6*, *E7*, *E5*, *E1*) across clusters, highlighting cluster 7 (terminally differentiated, low viral gene expression) and cluster 9 (productive amplification, all viral genes expressed). **l**, Volcano plot of cluster 9 versus all remaining clusters; genes significantly increased in cluster 9 are shown in red and those decreased in blue, with selected mitotic, kinetochore, histone and DNA-damage genes labelled. **m**, Gene ontology analysis of cluster 9 differential genes depicting pathways regulated by the genes expressed in cluster 9; dot size indicates gene count, and color indicates −log₁₀(*P*).

HPV genomes are amplified in differentiated cells that retain a G2/M program^6–8^, however neither feature alone is sufficient. Cycling basal-like cells that re-enter G2/M do not amplify and similar effects are seen in terminally differentiated cells that have left the cycle. UMAP embeddings of UD and D4 cells (**Fig. 1b**) resolved into ten clusters (**Fig. 1c**). Involucrin (*IVL*) was restricted almost entirely to cluster 7, identifying it as the most terminally differentiated population (**Fig. 1d,g**). *CDK1* and *TOP2A* were expressed across several clusters containing cells from both undifferentiated and differentiated conditions and were highest in cluster 9 (**Fig. 1e,f,h,i**), which was defined by a program of mitotic regulators, kinetochore and spindle components, differentiation markers and replication-dependent histones (**Fig. 1j**). Viral transcripts were detected across most clusters, as viral genomes are present in all cells (**Fig. 1k** and **Supplementary Fig. 2**). Clusters 2, 8 and 10 expressed *CDK1* and *TOP2A* at high levels but consisted predominantly of undifferentiated cells (**Fig. 1b,c,h,i**), while cluster 7 expressed differentiation markers but had exited the cycle and expressed viral genes poorly (**Fig. 1d,g,k**). Cluster 9 was the only population arising within the differentiated compartment that retained the full G2/M and DNA replication signature while expressing high levels of early viral genes required for amplification. This included *E1* and *E2*, which encode the origin-binding helicase and its loading factor, together with *E5*, *E6*, *E7* and the spliced *E1^E4* transcript (**Fig. 1k**). The difference between cluster 9 and the proliferative undifferentiated clusters is one of differentiation state rather than cell-cycle activity. At the same time the difference between cluster 9 and cluster 7 is one of G2/M cell-cycle activity rather than differentiation state.

Examining the expression of cellular genes in cluster 9 in comparison to the remaining clusters identified a set (**Fig. 1l**) enriched for the G2/M transition, DNA replication, DNA repair, viral carcinogenesis and DNA-damage-induced senescence (**Fig. 1m**). A protein–protein interaction network built from these genes resolved into a dense mitotic and kinetochore core along with a separate replication-dependent histone module, with TOP2A and NCAPD2 bridging the two (**Supplementary Fig. 3**). Among these genes we focused on *TOP2A* as it was highly expressed in cluster 9 and is engaged in the DNA transactions required to amplify a circular, chromatinized viral episome several thousand-fold. These activities include relief of the positive supercoiling generated ahead of replication forks, and separation of the interlinked daughter molecules produced when replication of a circular template terminates^26,27^. These studies identified a cluster of differentiated HPV positive cells using single cell RNA-seq analyses that expressed high levels of G2/M factors along with the topoisomerase, TOP2A, that was a prime candidate regulator of amplification and late gene expression.

### The cluster 9 program is consistent with a G2/M-restricted state that is retained in HPV-positive cells

We next wanted to confirm that the genes identified by single-cell RNA seq in the *TOP2A*-expressing cluster 9 are co-expressed within TOP2α-positive CIN612 cells in G2/M using a complementary FACS sorting method. For this analysis we included two additional genes: *NUSAP1*, a mitotic spindle-associated factor whose expression closely parallels that of *TOP2A*, and *H2AX*, a marker of DNA damage which is induced at high levels in cells undergoing amplification^9^ (**Fig. 2a,b**). FACS-based cell-cycle profiling by DNA content showed that upon differentiation uninfected primary human foreskin keratinocytes (HFKs) exhibited reduced numbers of G2/M cells (13.4% to 6.43%) as expected for normal cell-cycle exit. In contrast, HPV positive CIN612 cells exhibited increased numbers of cells in the G2/M fraction (6.90% to 9.92%) (**Fig. 2c**). Differentiating HPV-positive cultures therefore retain, and modestly expand, a G2/M population, which is where amplification is reported to occur^7,8^. Immunoblot analysis of FACS-sorted CIN612 cells revealed that TOP2α, NUSAP1, inhibitory pCDK1^Thr14/Tyr15^ and γH2AX began to increase in S phase and accumulated in G2/M. Importantly this accumulation increased further upon differentiation (**Fig. 2d,e**). Inhibitory phosphorylation of CDK1 at Thr14/Tyr15 marks cells arrested at the G2/M boundary rather than progressing through mitosis^7^, which indicates that differentiating HPV-positive cells are arrested in what we refer to as a pseudo-mitotic state. We observed similar results by co-immunofluorescence analysis, with nuclear TOP2α coincident with NUSAP1 (72.7% to 81.8% of TOP2α-positive nuclei in differentiated cells) and pCDK1 (85.2% to 89.7% in differentiated cells) in differentiated CIN612 but not HFK cells (**Fig. 2h,i**). In contrast, pS139-γH2AX increased significantly from 59.3% to 81.4% of TOP2α-positive nuclei (**Fig. 2i**). Taken together, cells expressing *TOP2A* also express the genes identified from the single-cell RNA sequencing analysis as important for viral genome amplification. Importantly we determined that TOP2α expression was confined to a subset of E1^E4-positive cells, and that this subset was substantially larger after 4 days of differentiation than in undifferentiated cultures (**Supplementary Fig. 4a**).

**Figure 2.**
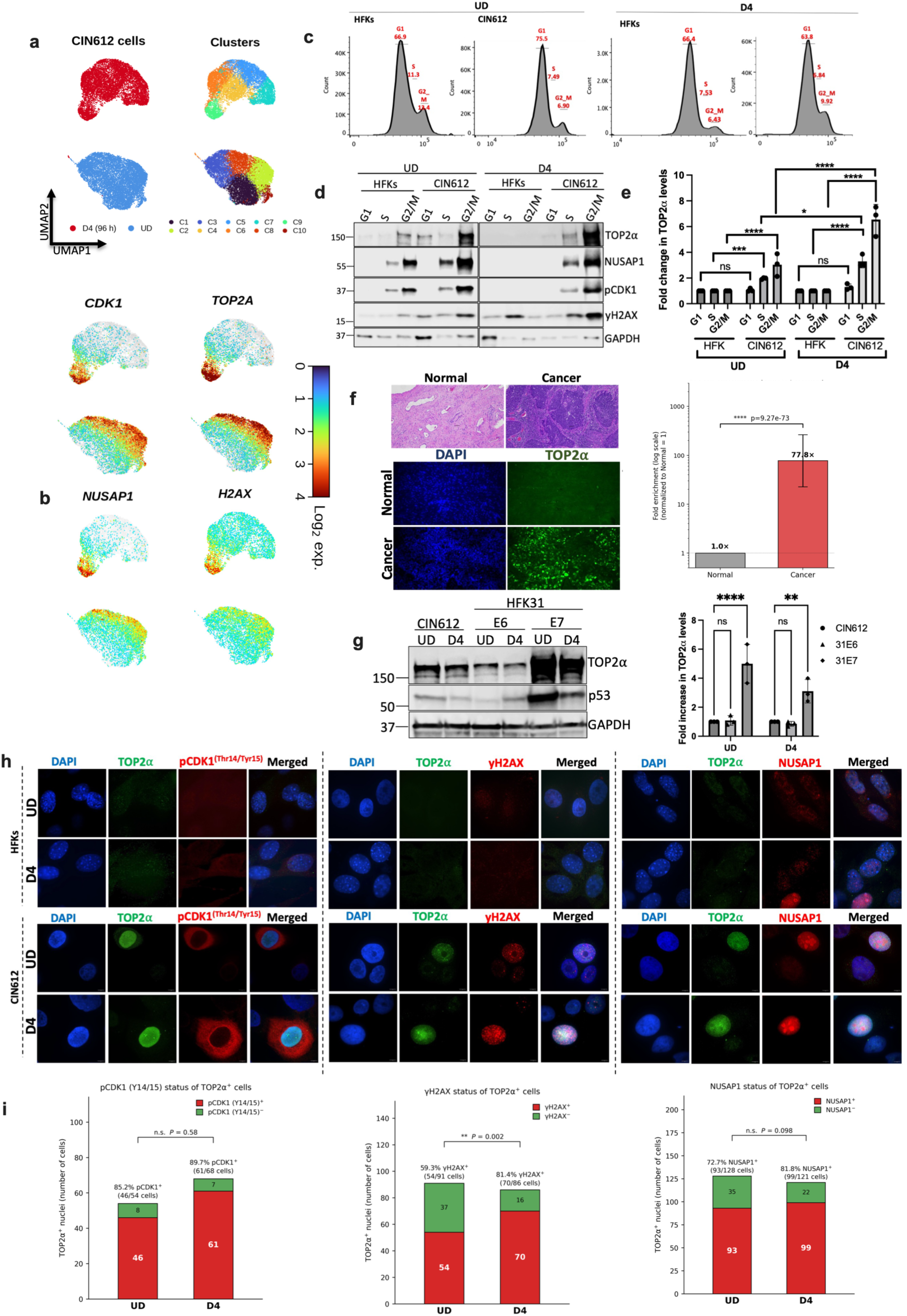
TOP2α accumulates with the G2/M programme in differentiating HPV-positive cells and is induced by E7. **a**, UMAP embeddings of CIN612 cells by condition and by cluster, with feature plots for *CDK1* and *TOP2A* (D4 upper, UD lower). **b**, Feature plots for *NUSAP1* and *H2AX* on the same embedding. **c**, Cell-cycle distribution determined by DNA content in undifferentiated (UD) and differentiated (D4) HFK and CIN612 cultures; percentages of cells in G1, S and G2/M are indicated. **d**, Immunoblot of FACS-sorted G1, S and G2/M fractions from UD and D4 HFK and CIN612 cultures probed for TOP2α, NUSAP1, pCDK1 (Thr14/Tyr15), γH2AX and GAPDH. **e**, Quantification of TOP2α levels from **d**, normalized to GAPDH (mean ± s.d., *n* = 3). **f**, Haematoxylin and eosin staining and TOP2α immunofluorescence of HPV-positive cancer biopsy tissue and matched normal epithelium on the same slide, with quantification of TOP2α enrichment (log scale, normalized to normal = 1). **g**, Immunoblot of CIN612 cells and HFKs expressing HPV31 E6 or E7 in UD and D4 states, probed for TOP2α, p53 and GAPDH, with quantification of TOP2α levels (mean ± s.d., *n* = 3). **h**, Co-immunofluorescence of UD and D4 HFK and CIN612 cells for DAPI, TOP2α and pCDK1 (Thr14/Tyr15), γH2AX or NUSAP1. **i**, Quantification of the CIN612 panels in **h**, showing the proportion of TOP2α-positive nuclei that are positive for pCDK1 (Thr14/Tyr15), γH2AX or NUSAP1 in UD and D4 cultures; cell numbers are indicated within each bar. Two-sided Fisher exact test. \**P* ≤ 0.05, \*\**P* ≤ 0.01, \*\*\**P* ≤ 0.001, \*\*\*\**P* ≤ 0.0001; ns, not significant.

We next examined whether elevated TOP2α is also a feature of HPV-associated disease *in vivo*. Immunofluorescence analysis of HPV-positive cancer biopsies showed that TOP2α was strongly elevated relative to normal epithelium present on the same slide (77.8-fold; **Fig. 2f**), consistent with the progressive increase in TOP2A reported across normal, low-grade, high-grade and carcinoma cervical tissue^33,34^. To determine which viral oncoprotein is responsible for this induction, TOP2α levels were compared in CIN612 cells and in HFKs stably expressing HPV31 E6 or E7. TOP2α was increased approximately 5-fold in undifferentiated and 3-fold in differentiated E7-expressing cells, whereas E6 had no significant effect (**Fig. 2g**). The same lysates were probed for p53, which was stabilized in the E7-expressing cells and reduced in the E6-expressing cells (**Fig. 2g**). Stabilization of p53 by E7 in the absence of E6 has been reported previously^35–37^, and the loss of p53 in the E6-expressing cells is the expected consequence of E6-directed degradation^16^, confirming that the E6 construct is active and that its lack of effect on TOP2α does not reflect a failure of expression. Together these data indicate that TOP2α is expressed at high levels both in differentiated cells undergoing productive replication and in cancers that often express only E6 and E7, and identify E7 as the viral determinant of its upregulation.

### TOP2α is confined to E4-positive cells and associates with amplifying genomes in stratified epithelium

It was next important to examine whether TOP2α is expressed in HPV E1^E4-expressing cells, using organotypic rafts that closely mimic the *in vivo* physiology of the epidermis and reproduce the complete viral life cycle. CIN612 and HFK organotypic rafts were grown at the air–liquid interface on collagen matrices seeded with NIH 3T3 J2 fibroblasts (**Fig. 3a**). Haematoxylin and eosin staining confirmed a fully stratified epithelium (**Fig. 3b**), and IF analysis revealed keratin 10 and loricrin were expressed in the suprabasal and granular compartments of the raft, establishing that differentiation proceeded in the presence of viral genomes (**Fig. 3c**).

**Figure 3.**
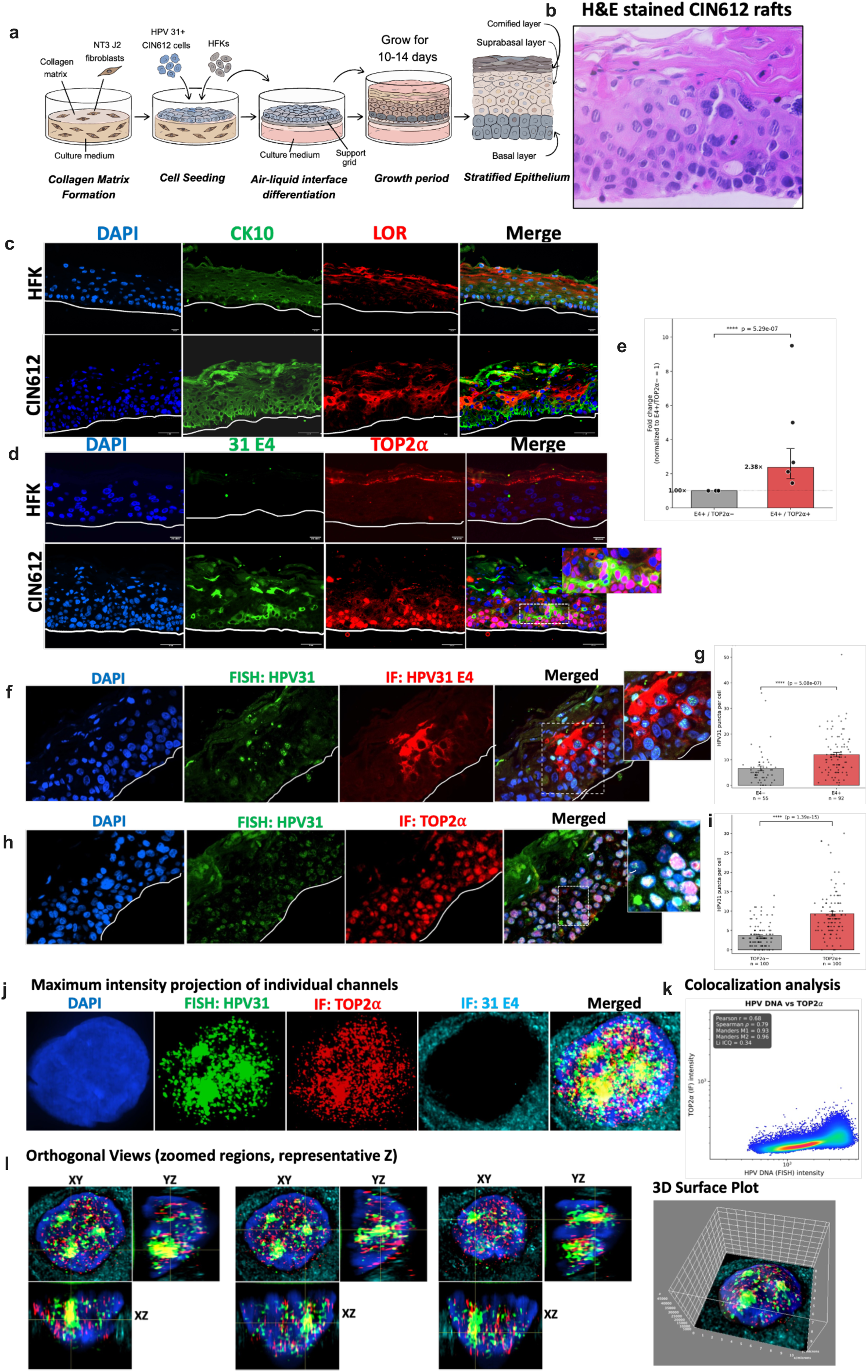
TOP2α is restricted to E1^E4-positive suprabasal cells and co-localizes with amplifying HPV31 genomes in organotypic raft culture. a, Scheme of organotypic raft culture. b, Haematoxylin and eosin staining of a CIN612 raft showing full stratification. c, Immunofluorescence for keratin 10 (CK10) and loricrin (LOR) in HFK and CIN612 rafts; the basement membrane is marked by a white line. d, Co-immunofluorescence for HPV31 E1^E4 (31 E4) and TOP2α in HFK and CIN612 rafts; the boxed region is enlarged in the inset. e, Quantification of d expressed as the fold change in TOP2α-positive relative to TOP2α-negative nuclei within the E4-positive compartment (mean ± s.e.m.). f, DNA FISH for HPV31 combined with immunofluorescence for HPV31 E1^E4. g, HPV31 puncta per nucleus in E4-negative (*n* = 55) and E4-positive (*n* = 92) cells. h, DNA FISH for HPV31 combined with immunofluorescence for TOP2α. i, HPV31 puncta per nucleus in TOP2α-negative and TOP2α-positive cells (*n* = 100 each). j, Maximum-intensity projections of individual channels (DAPI, HPV31 FISH, TOP2α IF, 31 E4 IF and merge) for a representative suprabasal nucleus. k, Colocalization analysis of HPV31 DNA and TOP2α intensities with Pearson, Spearman, Manders and Li ICQ coefficients. l, Orthogonal XY, YZ and XZ views of three representative nuclei and a 3D surface rendering. Two-sided Mann–Whitney *U* test; \*\*\*\**P* ≤ 0.0001.

To investigate whether TOP2α is confined to the cells amplifying HPV genomes, co-immunofluorescence for HPV31 E1^E4 and TOP2α was used to show nuclear TOP2α concentrated within E4-positive suprabasal cells of CIN612 rafts and absent from HFK rafts, in which TOP2α staining was weak and non-nuclear (**Fig. 3d**). Quantification showed that, within the E4-positive compartment, TOP2α-positive nuclei were 2.38-fold more frequent than TOP2α-negative nuclei (**Fig. 3e**).

It was next important to determine if the effects were specific to TOP2α or extended to other topoisomerases as previous studies have shown the involvement of TOP2β, TOP1 and TOP3β in viral replication in undifferentiated cells^28,29,38^. Interestingly, TOP1, TOP3β and TOP2β were each expressed throughout both the basal and suprabasal compartments of HFK and CIN612 rafts and showed no preferential association with E4-positive cells (**Supplementary Fig. 4c**). Similarly our single-cell data analysis also revealed that *TOP1*, *TOP3B* and *TOP2B* transcripts were distributed across all ten clusters, with no discrete enrichment in cluster 9, as seen in the case of *TOP2A*. This suggests that enrichment within the amplifying compartment is specific to TOP2α rather than to topoisomerases in general.

To investigate if TOP2α associates with amplified HPV DNA in differentiating cells, we combined DNA fluorescence *in situ* hybridisation (FISH) for HPV31 with immunofluorescence for E1^E4 and TOP2α in HPV-positive 3D organotypic raft cultures. Viral genomes formed large nuclear foci in suprabasal cells of raft cultures that coincided with both E1^E4 and TOP2α expression (**Fig. 3f,h**). The number of viral puncta per nucleus was significantly increased in E4-positive compared with E4-negative cells (**Fig. 3g**) as well as in TOP2α-positive compared with TOP2α-negative cells (**Fig. 3i**), confirming that both markers identify nuclei that are actively amplifying viral genomes.

High-resolution confocal imaging of HPV positive raft cultures further resolved their spatial relationship. Maximum-intensity projections showed HPV31 DNA and TOP2α occupying the same nuclear territory within E4-positive cells (**Fig. 3j**), and orthogonal reconstructions and surface rendering confirmed that this overlap persisted through the depth of the nucleus rather than arising from projection of separate signals (**Fig. 3l**). Line-intensity profiles showed coincident peaks (**Supplementary Fig. 4b**), and colocalization analysis gave Pearson *r* = 0.68, Spearman ρ = 0.79, Manders M1 = 0.93, M2 = 0.96 and Li ICQ = 0.34 (**Fig. 3k**), indicating that most of the TOP2α signal lies within nuclei that are HPV31 positive. Taken together, these observations demonstrate co-localization of TOP2α with HPV genomes undergoing amplification.

### TOP2α associates with the viral origin of replication and the early promoter

Having established that TOP2α is expressed preferentially in amplifying cells and co-localizes with HPV genomes, it was important to investigate how *TOP2A* expression is regulated and if it binds to HPV genomes. Western blot analyses indicate that total levels of TOP2α protein are significantly enriched in CIN612 cells compared to HFK cells (**Supplementary Fig. 5a**). Chromatin immunoprecipitation (ChIP) analysis^28^ revealed strong TOP2α enrichment at ALU repeats in CIN612 cells relative to HFKs, as expected for a host chromatin-associated enzyme that is itself elevated in HPV-positive cells, together with specific enrichment at the HPV31 upstream regulatory region (URR) in CIN612 cells (**Fig. 4a**). Mapping occupancy across the viral genome in UD and D4 cells revealed a differentiation-dependent increase at the URR (4.2-fold) and at the p97 early promoter (2.7-fold), with no significant change at the early or late polyadenylation sites (**Fig. 4b**). TOP2α is therefore preferentially recruited to the viral origin and early promoter at the point in the life cycle at which amplification is initiated, rather than distributed uniformly along the episome.

**Figure 4.**
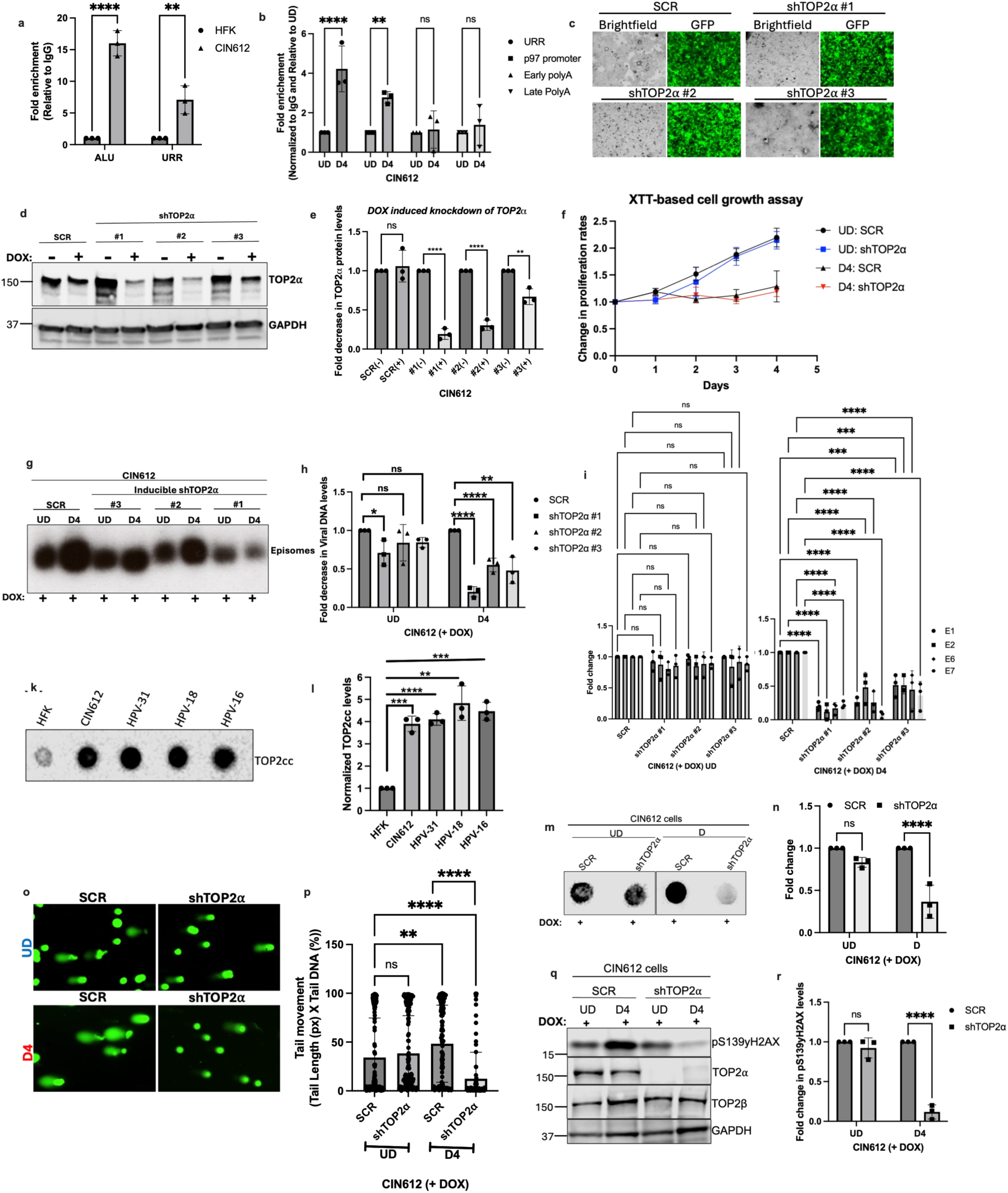
TOP2α binds the viral origin, is required for differentiation-dependent amplification, and generates cleavage complexes and DNA breaks. a, TOP2α chromatin immunoprecipitation in HFK and CIN612 cells at ALU repeats and the HPV31 URR, expressed as fold enrichment relative to IgG (mean ± s.d., *n* = 3). **b**, TOP2α occupancy across the HPV31 genome (URR, p97 promoter, early polyA, late polyA) in UD and D4 CIN612 cells, normalized to IgG and to UD. **c**, Brightfield and GFP images of doxycycline-inducible scrambled (SCR) and shTOP2α (#1–#3) CIN612 lines. **d**, Immunoblot for TOP2α and GAPDH in the inducible lines with and without doxycycline. **e**, Quantification of **d** (mean ± s.d., *n* = 3). **f**, XTT proliferation assay of SCR and shTOP2α CIN612 cultures in UD and D4 conditions. **g**, Southern blot of HPV31 episomes in doxycycline-treated SCR and inducible shTOP2α lines in UD and D4 conditions. **h**, Quantification of **g** (mean ± s.d., *n* = 3). **i**, RT– qPCR of *E1*, *E2*, *E6* and *E7* transcripts in doxycycline-treated UD and D4 cultures, normalized to GAPDH and to SCR. **k**, Dot blot of TOP2 cleavage complexes (TOP2cc) recovered by the RADAR assay from HFK, CIN612 and HFKs maintaining HPV31, HPV18 or HPV16 genomes. **l**, Quantification of **k** (mean ± s.d., *n* = 3). **m**, TOP2cc slot blot from doxycycline-treated SCR and shTOP2α CIN612 cultures in UD and differentiated conditions (labelled D in the panel). **n**, Quantification of **m**. **o**, Representative alkaline comet assay images from doxycycline-treated SCR and shTOP2α CIN612 cultures in UD and D4 conditions. **p**, Tail moment (tail length × tail DNA%) for the conditions in **o**; at least 100 nuclei per condition. **q**, Immunoblot of doxycycline-treated SCR and shTOP2α CIN612 cultures in UD and D4 conditions probed for pS139-γH2AX, TOP2α, TOP2β and GAPDH. **r**, Quantification of pS139-γH2AX from **q** (mean ± s.d., *n* = 3). Two-way ANOVA (panels **a**, **b**, **e**, **h**, **i**, **n**, **r**), one-way ANOVA with correction for multiple comparisons (panel **l**) or two-sided Mann–Whitney *U* test (panel **p**). \**P* ≤ 0.05, \*\**P* ≤ 0.01, \*\*\**P* ≤ 0.001, \*\*\*\**P* ≤ 0.0001; ns, not significant.

### TOP2α is required for differentiation-dependent genome amplification

To elucidate what effects depletion of TOP2α would have on HPV replication, CIN612 cells were generated that stably expressed shRNAs against *TOP2A*, using lentiviral expression vectors achieving approximately 90% depletion (**Supplementary Fig. 5a,b**) with only a modest effect on host cell proliferation (**Supplementary Fig. 5c**). Southern blot analysis showed a marked reduction in levels of HPV31 episomes (**Supplementary Fig. 5d**), which corresponded with reductions in *E1*, *E2*, *E6* and *E7* transcripts (**Supplementary Fig. 5e**).

Cells carrying a constitutive knockdown have undergone many divisions under selection, and loss of episomes in such a population cannot distinguish a requirement for the amplification step from cumulative failure of maintenance, or from outgrowth of cells that have adapted to TOP2α loss. We therefore generated three independent doxycycline-inducible, GFP-marked shTOP2α lines in the CIN612 background (**Fig. 4c**), each of which depleted TOP2α on treatment with doxycycline, by approximately 80%, 70% and 35% for shRNAs #1, #2 and #3 respectively (**Fig. 4d,e**). Proliferation was unaffected in UD cultures, and D4 cultures arrested regardless of whether the shRNA was expressed (**Fig. 4f**).

Acute depletion separated the two arms of the life cycle. Episomal HPV31 was largely unchanged in undifferentiated cells in which TOP2α was acutely reduced, with only shTOP2α #1 producing a small but significant decrease. In contrast, acute reduction of TOP2α with inducible shRNA vectors strongly reduced viral episomes in differentiated cells for all three shRNAs targeting *TOP2A* (**Fig. 4g,h**). Viral transcripts behaved identically, with *E1*, *E2*, *E6* and *E7* unchanged in UD cells and reduced by approximately 50–95%, depending on the shRNA, in differentiated D4 cells (**Fig. 4i**). Acute reduction of TOP2α in undifferentiated keratinocytes therefore has minimal short term effect on viral replication but results in a marked reduction in differentiation-dependent amplification.

Comparing the two systems, chronic depletion nearly eliminates the maintained episome pool whereas acute depletion does not. Both observations are internally consistent if maintenance carries a low-level requirement for TOP2α that becomes apparent only after many rounds of replication, whereas amplification requires it acutely.

### TOP2α forms cleavage complexes and generates programmed breaks on the viral genome

TOP2α relieves topological stress by cleaving both strands of DNA, passing a second duplex through the break and resealing it, transiently forming a covalent enzyme–DNA intermediate termed the TOP2 cleavage complex (TOP2cc)^26,27^. If TOP2α acts directly on viral DNA during amplification, TOP2cc should be elevated in HPV-positive cells and should depend on differentiation.

To investigate the contribution of these TOP2cc intermediates to viral functions, TOP2cc were recovered by chaotropic lysis and nucleic-acid precipitation (the RADAR assay, rapid approach to DNA adduct recovery) and detected by dot blot^39,40^. This showed approximately 4–5-fold higher TOP2cc levels in CIN612 cells and in HFKs carrying HPV31, HPV18 or HPV16 genomes than in HFKs alone (**Fig. 4k,l**). These results indicate that elevated cleavage complex formation occurs in different high-risk HPV types rather than in a single genotype or cell line. TOP2cc levels were reduced by approximately 75% in stable shTOP2α cells (**Supplementary Fig. 6a,b**). Using the inducible system, levels were unchanged in UD but decreased by over 50% in differentiated cells (**Fig. 4m,n**), consistent with the amplification phenotype.

Cleavage complexes that are not resealed give rise to protein-linked double-strand breaks^27,41^. Alkaline comet assays measuring DNA breaks showed that differentiation increased tail moment in control CIN612 cells and that inducible TOP2α depletion reduced it below the undifferentiated level in D4 cells (**Fig. 4o,p**). Stable knockdown of TOP2α gave a similar result (**Supplementary Fig. 6c,d**). γH2AX followed a similar pattern, decreasing by approximately 65% in stable knockdown cells (**Supplementary Fig. 6e,f**), and by approximately 90% in differentiated inducible cells (**Fig. 4q,r**). To ensure that TOP2α did not have an indirect effect on TOP2β to induce increases in TOP2cc, TOP2β levels were examined by western blot and found to be unchanged (**Fig. 4q** and **Supplementary Fig. 6e**). Furthermore, studies in raft cultures showed TOP2β is uniformly expressed across the epithelium (**Supplementary Fig. 4c**), suggesting that the two type II enzymes have non-redundant functions in HPV replication and maintenance.

### TOP2α maintains the G2/M program, and E7 induces TOP2α through FOXM1

It was next important to determine whether the requirement for TOP2α is limited to effects on DNA amplification and late gene expression or if it had a general effect in regulating entry into G2/M. G2/M fractions from doxycycline-treated inducible shTOP2α and control cultures in UD and D4 differentiated states were isolated by FACS and profiled by bulk RNA sequencing. In UD G2/M cells, gene-set enrichment analysis of TOP2α-depleted CIN612 cells showed reduced E2F target genes, G2/M checkpoint and MYC target signatures, with the induction of TNFα/NF-κB, p53, cholesterol homeostasis, myogenesis, androgen-response, hypoxia and apoptosis pathways (**Fig. 5a,b**). Effects were substantially more extensive in D4 differentiated G2/M cells, where E2F targets, the G2/M checkpoint, the mitotic spindle and DNA repair were all depleted, together with interferon-α, interferon-γ, IL-2–STAT5, mTORC1, oxidative phosphorylation, epithelial–mesenchymal transition and estrogen-response signatures, while p53 and unfolded protein responses were induced (**Fig. 5c,d**). Cluster 9 genes, including *UBE2C*, *CDC20*, *TPX2*, *AURKA*, *AURKB*, *PLK1*, *CENPE*, *CENPF*, *BUB1*, *NUSAP1* and *CDK1*, were among the most strongly altered transcripts, consistent with those identified in the TOP2α knockdowns. Interestingly, the interferon signatures are not specifically enriched in cluster 9.

**Figure 5.**
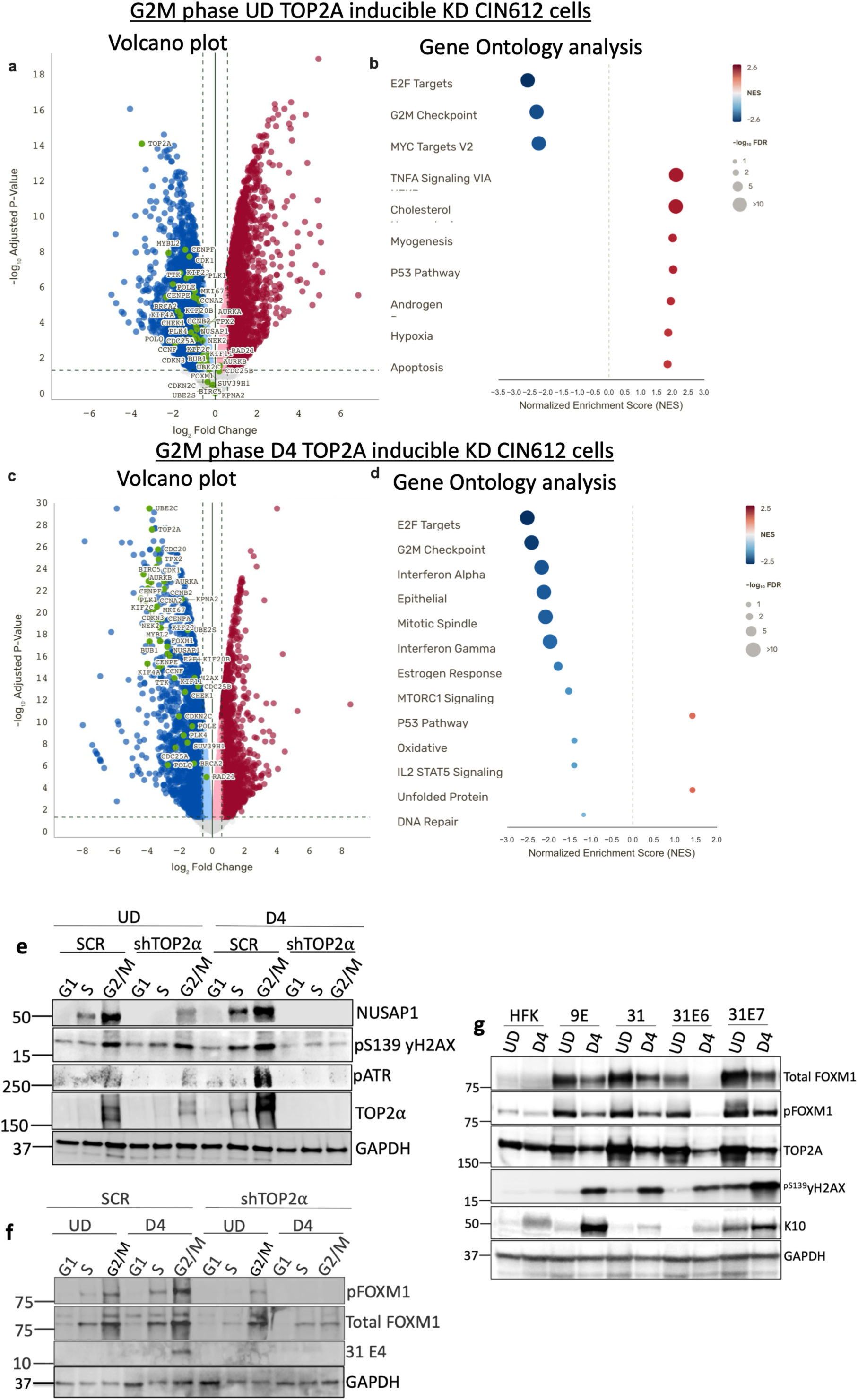
TOP2α sustains the G2/M transcriptional programme and FOXM1 activation, and E7 induces TOP2α and FOXM1. **a**, Volcano plot of differentially expressed genes in FACS-purified G2/M cells from undifferentiated doxycycline-treated inducible shTOP2α versus scrambled control CIN612 cultures; cluster 9 genes are highlighted in green. **b**, Hallmark gene-set enrichment analysis of the comparison in **a**; dot colour indicates normalized enrichment score (NES) and size indicates −log₁₀ FDR. [The panel is titled “Gene Ontology analysis” in the artwork but shows MSigDB Hallmark sets; please retitle.] **c**, Volcano plot for the same comparison in differentiated (D4) G2/M cells. **d**, Gene-set enrichment analysis of the comparison in **c**. **e**, Immunoblot of FACS-sorted G1, S and G2/M fractions from UD and D4 scrambled and shTOP2α CIN612 cultures probed for NUSAP1, pS139-γH2AX, pATR, TOP2α and GAPDH. **f**, Immunoblot of the same sorted fractions probed for pFOXM1, total FOXM1, HPV31 E1^E4 (31 E4) and GAPDH. [The two UD blocks in this panel are labelled G1 and S only, although twelve lanes are run; please label the third lane of each UD block or state in the legend that the UD G2/M fractions were not recovered.] **g**, Immunoblot of UD and D4 lysates from HFK, CIN612 9E, HFK31, and HFKs expressing HPV31 E6 (31E6) or E7 (31E7), probed for total FOXM1, pFOXM1, TOP2A, pS139-γH2AX, keratin 10 (K10) and GAPDH.

Western blot analysis of the same sorted fractions confirmed that the observations from the RNA-seq data were present at the protein level. NUSAP1 and γH2AX accumulated in the G2/M fraction of differentiated control cells and were lost upon TOP2α depletion, most markedly at D4; phosphorylated ATR followed the same pattern, consistent with loss of the DNA-damage signalling that accompanies amplification (**Fig. 5e**). FOXM1 has been reported to control expression of the G2/M gene cluster^30,31^ and is recruited to the promoters of these genes through CHR elements by the MuvB–B-MYB (MMB) complex^32,42–44^. Genome-wide occupancy studies have reported FOXM1 at the promoters of *PLK1*, *AURKB* and *CCNB1*^45^, and ChIP together with reporter assays confirm direct activation of *TOP2A*^46^. The genes that define cluster 9 — *CENPF*, *NEK2*, *PLK1*, *AURKB*, *CDC25C*, *CCNB2*, *UBE2C*, *NUSAP1*, *CENPE*, *BUB1*, *CDCA8* and *TOP2A* — correspond closely to this characterized FOXM1 target set^30,31,45,46^, and loss of FOXM1 produces the mitotic and chromosome-segregation defects expected from coordinate loss of these genes^47^. Since FOXM1 activity is controlled by phosphorylation at the G2/M transition rather than by abundance alone^48,49^, the same fractions were probed for phosphorylated FOXM1 (pFOXM1) and total FOXM1. Both were increased in the G2/M fraction upon differentiation, in parallel with increases in levels of NUSAP1 and γH2AX, placing FOXM1 activation in the same fraction in which TOP2α accumulates (**Fig. 2d,e**) and in which amplification occurs. Upon depletion of TOP2α, pFOXM1 and total FOXM1 were reduced in differentiated cells and less so in undifferentiated cells (**Fig. 5f**). These observations place TOP2α within an auto-regulatory circuit with FOXM1 to control expression of G2/M genes and amplification.

It was next important to correlate these changes with HPV late gene expression. The same fractions that were depleted for TOP2α were probed for the late E1^E4 protein, which marks cells that have entered the productive program^19,20^. E1^E4 was increased in the G2/M fraction of CIN612 cells upon differentiation and was markedly decreased in differentiated G2/M cells upon depletion of TOP2α (**Fig. 5f**). These studies indicate that the requirement for TOP2α extends beyond amplification to the entire productive program. Importantly, these data also show that the G2/M cluster 9 program is not sustained in the absence of TOP2α.

It was next useful to determine which viral gene is responsible for the increased expression of *TOP2A*, and how that expression is driven. Western blot analysis was performed on HFKs, HFKs stably maintaining HPV31 genomes, HFKs expressing HPV31 E6 or E7, and CIN612 9E cells following retroviral infection. Increases in the protein levels of TOP2α, total FOXM1 and pFOXM1 were observed in E7 expressing cells, whereas E6 had little effect (**Fig. 5g**). A modest decrease in pFOXM1 and total FOXM1 was observed in whole-cell lysates upon differentiation of the E7-expressing cells, in contrast to their increase in the sorted G2/M fraction. The same lysates were also probed for pS139-γH2AX, which was elevated in the E7-expressing cells (**Fig. 5g**). Accumulation of γH2AX is expected since expression of E7 alone is sufficient to activate ATM and its downstream targets and to induce DNA breaks^9,50^.

Taken together, these studies define a differentiation-restricted, TOP2α-dependent state that licenses productive HPV replication. E7 drives the accumulation and activation of FOXM1, which in turn sustains expression of the cluster 9 G2/M programme including *TOP2A*; TOP2α is then required both for entry into G2/M and for the DNA transactions and programmed breaks that accompany genome amplification, and its loss collapses the FOXM1 programme and abolishes late gene expression.

## Discussion

Productive HPV replication is dependent upon epithelial differentiation but is restricted to a subset of cells that re-enter G2/M despite the presence of viral genomes in most cells. The requirement for G2/M entry reflects the dependency of HPV genome amplification on activation of the homologous DNA repair pathways, ATM and ATR. Our studies have determined that TOP2α is a prime regulator of productive viral replication through its control of entry into G2/M. Profiling differentiating HPV31-positive keratinocytes with single-cell RNA-seq identified a single differentiated population, cluster 9, in which the host and viral requirements for amplification coincide. TOP2α was highly expressed in these cells and knockdown specifically in differentiating cells blocked G2/M entry, amplification and late viral gene expression.

Ten clusters of undifferentiated and differentiated HPV positive cells were identified by single cell analysis, however, only one cluster, cluster 9, was defined by high levels of late viral transcripts for E1^E4 and G2/M entry. Viral transcripts were detected across most differentiated clusters, as all cells contain HPV genomes; however only cluster 9 expressed late genes at high levels and had re-entered G2/M. In addition, proliferation ability alone was also not sufficient to activate late viral functions as the most proliferative populations are undifferentiated clusters 2, 8 and 10, all of which express high levels of *CDK1* and *TOP2A*, but failed to induce late viral functions. It appears that cluster 9 is unique in that it arises within the differentiated compartment while retaining a complete G2/M and DNA-replication signature as well as expressing the viral replication genes.

Interestingly, while stable expression of shRNAs to *TOP2A* impairs viral replication in undifferentiated cells, acute, inducible depletion leaves episome copy number and viral transcription largely intact. In contrast, acute depletion in differentiating cells blocks viral genome amplification, entry into G2/M and late gene activation. This suggests that TOP2α is required for the amplification step itself and that its loss also destabilizes the expression of G2/M and E2F genes in these cells.

Our studies indicate that TOP2α is the prime topoisomerase important for transition into G2/M in differentiating HPV positive cells as it is the only one whose levels increase and whose knockdown blocks entry. At least a portion of the TOP2α effect is due to transcriptional effects as E2F target genes, G2/M checkpoint and epithelial– mesenchymal transition signatures are all affected by TOP2α knockdown. Previous studies showed that knockdown of TOP1 and TOP3β in undifferentiated cells also altered expression but in pathways that are distinct from that seen with depletion of *TOP2A* in differentiating cells^29^. This may be due to binding to specific sets of genes or altered expression of common transcription factors like FOXM1. Topoisomerases also regulate R-loop formation, and these structures have been shown to be critical for HPV late functions^13,27,51,52^. Further, TOP2α may have specific enzymatic activities that are particularly suited to HPV genome amplification. Amplification requires that a covalently closed circular, chromatinized template be replicated several thousand-fold within a differentiated cell. Both the positive supercoiling generated ahead of converging forks and the catenanes produced when replication of a circular template terminates require a type II activity for resolution; type I enzymes relax supercoils but cannot decatenate covalently closed duplex circles^26,27,53^. The differentiation-dependent enrichment of TOP2α at the URR and the p97 promoter, rather than uniform distribution along the episome, is consistent with recruitment at the point of replication initiation. HPV amplification has been suggested to occur either by a rolling circle mechanism or random priming^6^ and TOP2α may also play a role in these forms of replication, but these have not been fully elucidated.

HPV-positive cells induce high levels of DNA breaks which are required to activate the ATM and ATR pathways on which amplification depends^9,10,13,50^. Our studies indicate that TOP2α induces high levels of DNA breaks in differentiating cells undergoing amplification and that TOP2cc levels are elevated across multiple high-risk HPV positive cells. The levels of these breaks increase substantially on differentiation, and TOP2α depletion reduces both the TOP2cc complexes and over half of the DNA breaks. This suggests that a substantial part of the DNA damage that HPV-positive cells sustain is generated by the enzymatic action of TOP2α. Importantly, the levels of γH2AX and DNA breaks are reduced on TOP2α knockdown, which is the result of inhibition of break formation and not because damage has been repaired. E7 is the oncoprotein responsible for activating ATM, and this may occur in part through the induction of TOP2α via FOXM1.

E7 has been reported to associate with the FOXM1 activator in a manner that requires its LXCXE motif, in addition to degrading Rb^14,17,54^. FOXM1 has been reported to be the transcriptional driver of G2/M genes including those expressed in cluster 9^30–32,45,48^. FOXM1 is a transcription factor whose regulon includes *TOP2A*^46^. E7 has been reported to cooperate with FOXM1 to activate mitotic genes^54^, and *FOXM1* is among the genes most strongly repressed when E7 is silenced in HPV-positive cells, in an Rb-dependent manner^55^. FOXM1 is itself over-expressed in cervical cancer and its expression correlates with disease progression^56^. Our studies indicate that TOP2α and FOXM1 may be in an auto-regulatory loop controlling G2/M entry.

Acute depletion of TOP2α lowers activated and total FOXM1 in the differentiated G2/M fraction while leaving undifferentiated cells largely unaffected, and the same depletion reduces *E6* and *E7* transcripts in differentiated cells. Since E7 is itself required to maintain FOXM1^54,55^, one explanation is that there is an auto-regulated circuit in which FOXM1 regulates TOP2α expression, TOP2α supports amplification and viral transcription, which results in increased levels of E7, which in turn sustains FOXM1.

The observation that activated FOXM1 is reduced upon TOP2α depletion specifically in differentiated cells, when viral gene expression is also lost, is consistent with such a feedback arm. These studies identify TOP2α as a critical regulator of differentiation-dependent HPV amplification by acting to control G2/M entry and viral replication working in a feedback loop with FOXM1.

## Materials and Methods

### Cell culture and differentiation

HPV31-positive CIN612 9E keratinocytes, derived from a cervical intraepithelial neoplasia grade 1 biopsy and maintaining HPV31b as episomes^22,23^, were grown in E-medium supplemented with epidermal growth factor in the presence of mitomycin C-treated NIH 3T3 J2 fibroblast feeders, as described^21,24^. Primary human foreskin keratinocytes (HFKs) were isolated from de-identified neonatal foreskin tissue and maintained under the same conditions. HFK lines stably maintaining recircularized HPV16, HPV18 or HPV31 genomes (HFK16, HFK18, HFK31) were generated by co-transfection of religated viral genomes with a neomycin-resistance plasmid followed by G418 selection^24,57^. HFKs expressing HPV31 E6 or E7 were generated by retroviral transduction with pLXSN-based constructs and selected in G418^58^. Differentiation was induced by calcium switch: cells were seeded without feeders and transferred sequentially to medium containing 0.07 mM Ca²⁺ (24 h), low 0.03 mM Ca²⁺ (24 h) and high 1.5 mM Ca²⁺ (96 h), with harvest at 144 h^23,25^. Differentiation was confirmed by keratin 10 immunoblot and by induction of the spliced *E1^E4* transcript by RT-PCR. Cell viability was determined by trypan blue exclusion.

### Organotypic raft culture

Organotypic rafts were prepared essentially as previously described^21,24^. Collagen matrices seeded with NIH 3T3 J2 fibroblasts were overlaid with CIN612 or HFK cells, raised to the air–liquid interface on stainless steel grids and grown for 10–14 days with E-medium changed every other day. Rafts were fixed in 4% paraformaldehyde for 48 h at 4 °C, embedded in paraffin and sectioned at 4 µm. Sections were stained with haematoxylin and eosin or processed for immunofluorescence and DNA FISH.

### shRNA-mediated knockdown

Stable knockdown used lentiviral pLKO.1 vectors encoding shRNAs against *TOP2A* or a scrambled (SCR) control, with puromycin selection. Doxycycline-inducible knockdown used GFP-marked Tet-On lentiviral vectors carrying three independent *TOP2A* shRNAs (#1, #2, #3) (Dharmacon, #V3SH11255-01EG7153) or a scrambled control (Dharmacon, #VSC11651). Knockdown was induced with doxycycline, maintained throughout the differentiation time course, and verified by immunoblot at each harvest point. Transduction efficiency was monitored by GFP fluorescence.

### Single-cell RNA sequencing and analysis

Undifferentiated (UD) and day four differentiated (D4) CIN612 cultures were dissociated to single-cell suspensions and processed for 10x Genomics single-cell RNA sequencing. Libraries were generated by Northwestern Single Cell Core Facility and sequenced by Admira. Reads were aligned with Cell Ranger to a custom reference combining GRCh38 with the complete HPV31 genome, with each viral open reading frame annotated as an individual feature. Because standard barcode filtering preferentially removes viral reads in this system, reads assigned to low-confidence barcodes were retained in the count matrix.

### Cell-cycle analysis, sorting and bulk RNA sequencing

Cells were fixed, stained for DNA content and analysed by flow cytometry; G1, S and G2/M fractions were sorted for immunoblot analysis. For transcriptional profiling, G2/M fractions were sorted from doxycycline-treated inducible shTOP2α and scrambled control cultures in UD and D4 states and processed for bulk RNA sequencing using Plasmidosaurus.

### Western blot analysis

Cells were harvested and lysed in sample lysis buffer. Lysates were resolved by SDS–PAGE, transferred to nitrocellulose and probed with antibodies against anti-rabbit TOP2α (1:500, CST), anti-rabbit TOP2β (1:500, CST), anti-rabbit FOXM1 (1:500, CST), phosphorylated FOXM1 (pT600FOXM1; 1:500, CST), anti-rabbit NUSAP1 (1:2000,, anti-rabbitpCDK1 (Thr14/Tyr15, 1:500, CST), Anti-rabbit pATR (1:500, CST), anti-rabbit pSer139-γH2AX (1:3000, CST), anti-mouse p53 (1:4000, Santa Cruz), anti-mouse cytokeratin 10 (1:5000, Santa Cruz) and anti-mouse GAPDH (1:6000, Santa Cruz) followed by respective secondary HRP-conjugated antibodies (1:3000, CST). Signals were detected by enhanced chemiluminescence, quantified in ImageJ and normalized to GAPDH.

### Immunofluorescence and quantification

Monolayer cultures and raft sections were fixed, permeabilized, blocked and incubated with primary antibodies followed by fluorophore-conjugated secondary antibodies, and counterstained with DAPI. Images were acquired by confocal microscopy with identical settings across conditions within an experiment. For co-positivity scoring, nuclei were segmented on the DAPI channel and per-nucleus mean intensities were thresholded by the Otsu method, set independently for each image after linear unmixing and background subtraction; only within-image fractions were compared. Enrichment of TOP2α in E4-positive relative to E4-negative cells was computed on segmented nuclei from matched fields.

### DNA FISH combined with immunofluorescence

Raft sections were dewaxed, rehydrated, subjected to antigen retrieval and hybridized with a labelled HPV31 whole-genome probe. Following hybridization and stringency washes, sections were processed for immunofluorescence against HPV31 E4 or TOP2α and counterstained with DAPI. Confocal z-stacks were acquired for maximum-intensity projection, orthogonal reconstruction and surface rendering. Colocalization was computed on nuclear regions of interest after background subtraction, reporting Pearson and Spearman correlation coefficients, Manders coefficients M1 and M2, and the Li intensity correlation quotient. Puncta per nucleus were counted on segmented nuclei and compared by two-sided Mann–Whitney *U* test.

### Southern blotting

Total cellular DNA was isolated^9,24^ and separated on 1% agarose gels, transferred to nylon membranes and hybridized with a ³²P-labelled full-length HPV31 genomic probe. Signals were quantified by using and normalized to total DNA loaded.

### RT–qPCR

Total RNA was extracted, and reverse transcribed to generate cDNA. Viral transcripts (*E1*, *E2*, *E6*, *E7* and spliced *E1^E4*) and *TOP2A* were quantified by SYBR-based qPCR and normalized to GAPDH, with fold changes calculated by the ΔΔCt method relative to the stated control. Primer sequences are listed previously^29^.

### Chromatin immunoprecipitation

ChIP was performed as described for topoisomerases in HPV-positive keratinocytes^28,29^. Cells were crosslinked in formaldehyde, quenched, lysed and sonicated. Chromatin was immunoprecipitated with an antibody against TOP2α or with matched IgG, washed, eluted and reverse crosslinked. Recovered DNA was quantified by qPCR with primer pairs spanning the HPV31 URR, the p97 early promoter, and the early and late polyadenylation regions; ALU repeats served as a positive control for host chromatin. Enrichment is expressed relative to IgG.

### TOP2 cleavage complex (RADAR) assay

TOP2cc were recovered by the RADAR (rapid approach to DNA adduct recovery) procedure^39,40^. Cells were lysed in guanidinium-based buffer, DNA with covalently bound protein was precipitated in ethanol, washed and resuspended, and equal quantities of DNA were applied to nitrocellulose by dot blot. Membranes were probed with an antibody to TOP2α and developed by enhanced chemiluminescence. Signals were normalized to DNA loading, determined in parallel.

### Alkaline comet assay

Cells were embedded in low-melting-point agarose on slides, lysed, unwound under alkaline conditions and electrophoresed. Slides were neutralized, stained with a DNA-binding fluorophore and imaged. Tail moment was calculated as tail length (px) × tail DNA (%). At least 100 nuclei were scored per condition per replicate using FIJI COMET plugin.

### XTT-based Cell proliferation assay

Cell viability and growth pattern were assessed using the XTT colorimetric assay. SCR control CIN612 cells and topoisomerase knockdown cell lines including stable and inducible TOP2A knockdown, were seeded at 5,000 cells per well in 96-well plates in triplicate. Cell viability was measured at Days 1, 2, 3, 4, 6, and 8 post seeding for stable knockdown cells and at Days 1, 2 3, and 4 for inducible knockdown cells by adding 50 μL of XTT reagent per well and incubating for 4 h at 37 °C (as per the manufacturer’s protocol, Thermo Scientific). Absorbance was measured at 450 nm with 630 nm reference wavelength using a microplate reader. Growth curves were plotted using GraphPad Prism software, with all values normalized to SCR control to determine relative cell viability and proliferation rates.

### Statistics and reproducibility

Unless stated otherwise, data are presented as mean ± s.d. from three biologically independent experiments. Statistical analyses were performed in GraphPad Prism. Comparisons between two groups used two-sided unpaired Student *t*-tests. Comparisons across two factors — cell-cycle fraction and differentiation state, or genotype and differentiation state — used two-way ANOVA with correction for multiple comparisons. Per-nucleus distributions from the comet and FISH experiments, which are not normally distributed, were compared by two-sided Mann–Whitney *U* test, and co-positivity proportions by two-sided Fisher exact test. Significance is indicated as \**P* ≤ 0.05, \*\**P* ≤ 0.01, \*\*\**P* ≤ 0.001, \*\*\*\**P* ≤ 0.0001; ns, not significant.

### Data availability

Single-cell and bulk RNA sequencing data sets will be deposited in the NCBI Gene Expression Omnibus (GEO) and accession numbers will be provided prior to publication.

## Acknowledgements

This work was supported by grants to L.A.L. from the National Cancer Institute R01CA1422861 and R01CA295739. Imaging was performed on a Nikon SoRa system (purchased with the support of 1S10OD032270-01) at Northwestern University’s Center for Advanced Microscopy (RRID: SCR_020996), generously supported by NCI CCSG P30 CA060553 awarded to the Robert H Lurie Comprehensive Cancer Center. This research was supported in part by resources provided by the Northwestern University Skin Biology and Diseases Resource-based Center (P30AR075049), Chicago, IL with support from the NIH/NIAMS. Any opinions, findings, and conclusions or recommendations expressed in this material are those of the author(s) and do not necessarily reflect the views of the Northwestern University Skin Biology and Diseases Resource-based Center or the NIH/NIAMS. We would like to thank Dr. Piyush Garg, Lead AI Scientist at Point 72 who helped us with identifying HPV genes transcripts in our single cell RNA sequencing analysis.

## Supplementary information

All supplementary information is provided as a separate file.

