## Supplemental Figures 1-6 for "Single-cell analysis identifies TOP2α as a critical regulator of G2/M entry and differentiation-dependent productive HPV replication"

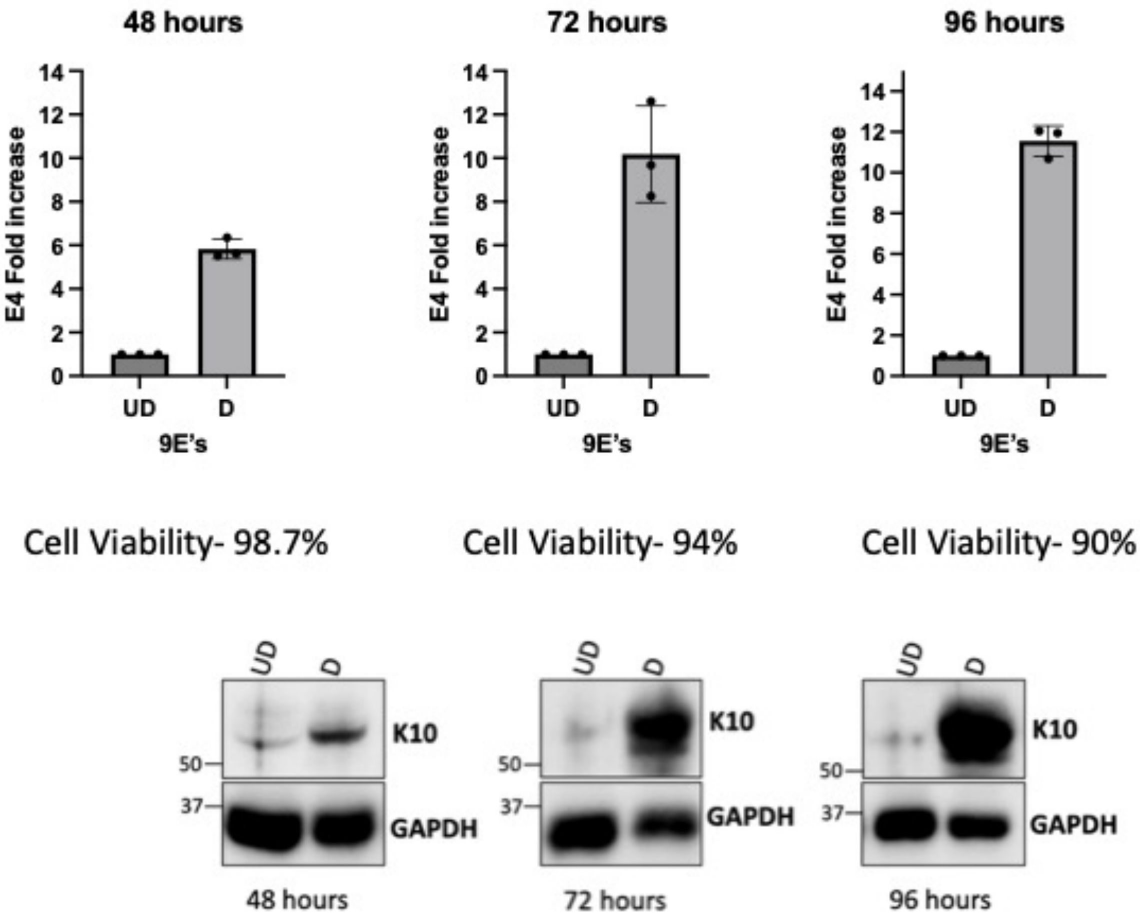


**Supplementary Figure 1 | Validation of the calcium-induced differentiation time course.** RT–qPCR of the spliced *E1^E4* transcript in undifferentiated (UD) and differentiated (D) CIN612 9E cultures at 48, 72 and 96 h after addition of 1.5 mM calcium (mean ± s.d., $n = 3$), with the corresponding cell viability determined by trypan blue exclusion indicated above each panel. Lower panels, immunoblot for keratin 10 (K10) and GAPDH at each time point. The 96 h time point (144 h total) was selected for single-cell RNA sequencing.

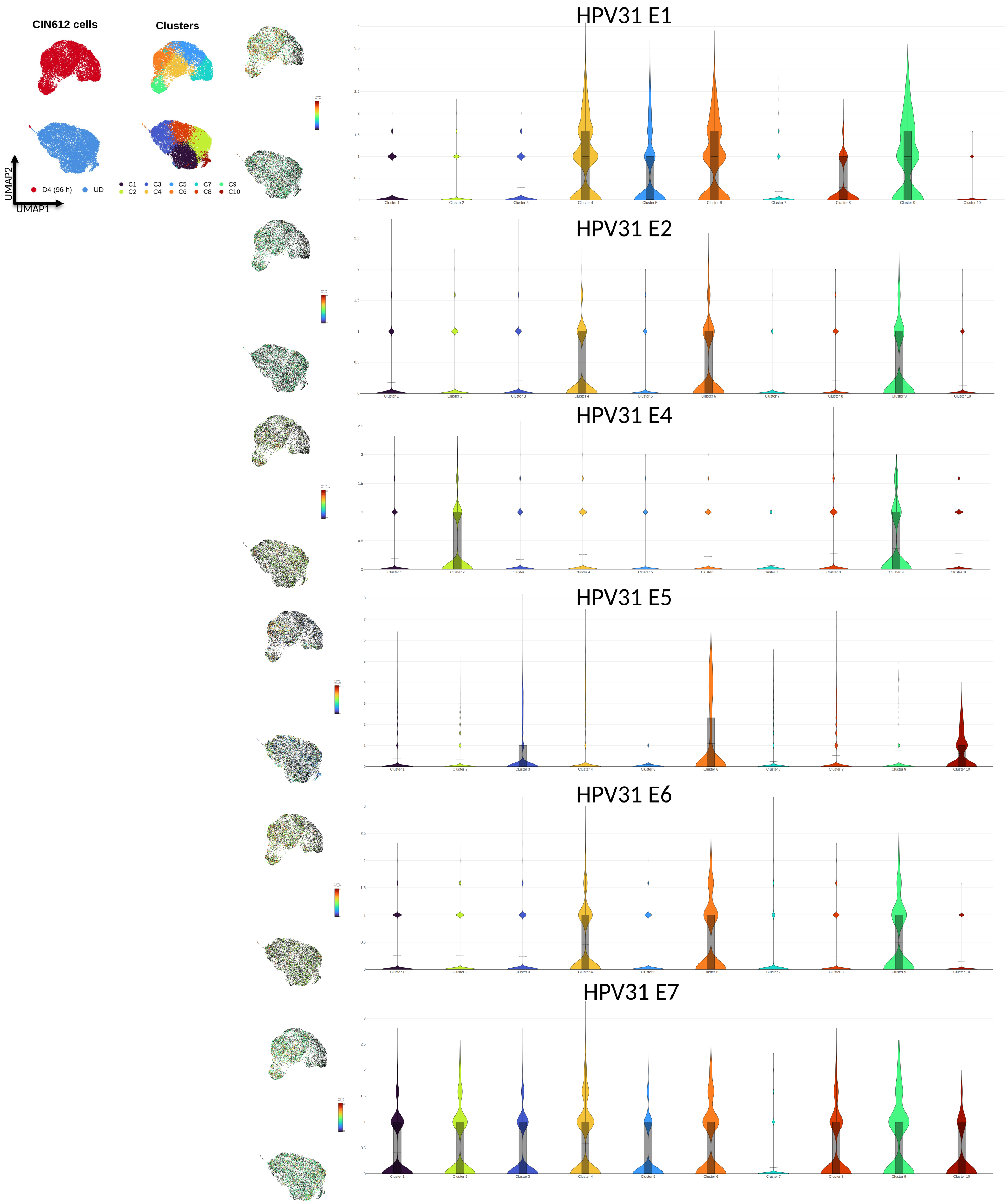


**Supplementary Figure 2 | Expression of individual HPV31 genes across the ten clusters.** Reference UMAP embeddings coloured by condition and by cluster (left), with feature plots and per-cluster violin plots of log-normalized expression for individual HPV31 gene.

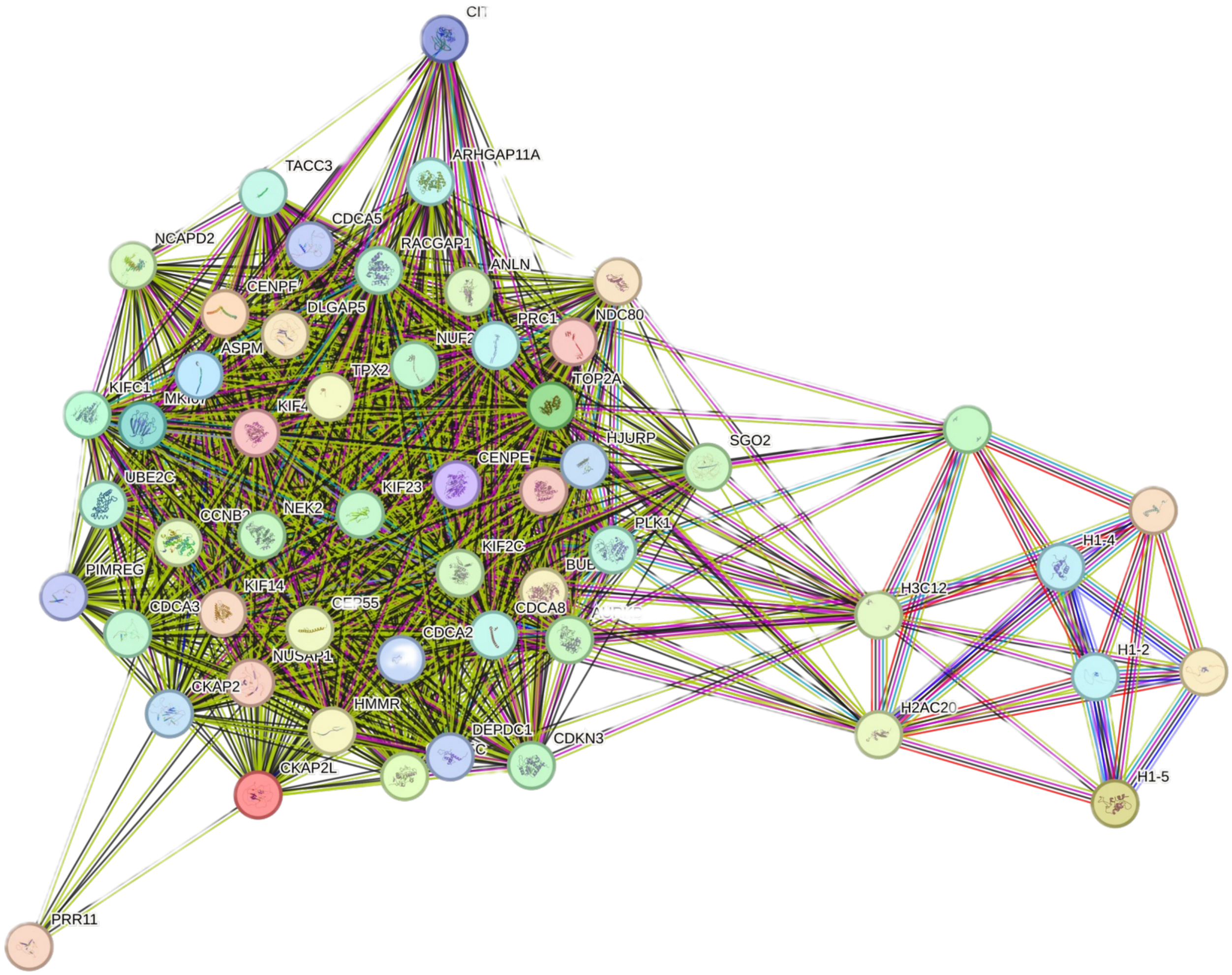


**Supplementary Figure 3 | Protein–protein interaction network of cluster 9 marker genes.** STRING network built from the top cluster 9 marker genes and visualized in Cytoscape. The network resolves into a dense mitotic and kinetochore module (left) and a separate replication-dependent histone module (right, H3C12, H2AC20, H1-2, H1-4, H1-5), with TOP2A and NCAPD2 bridging the two.

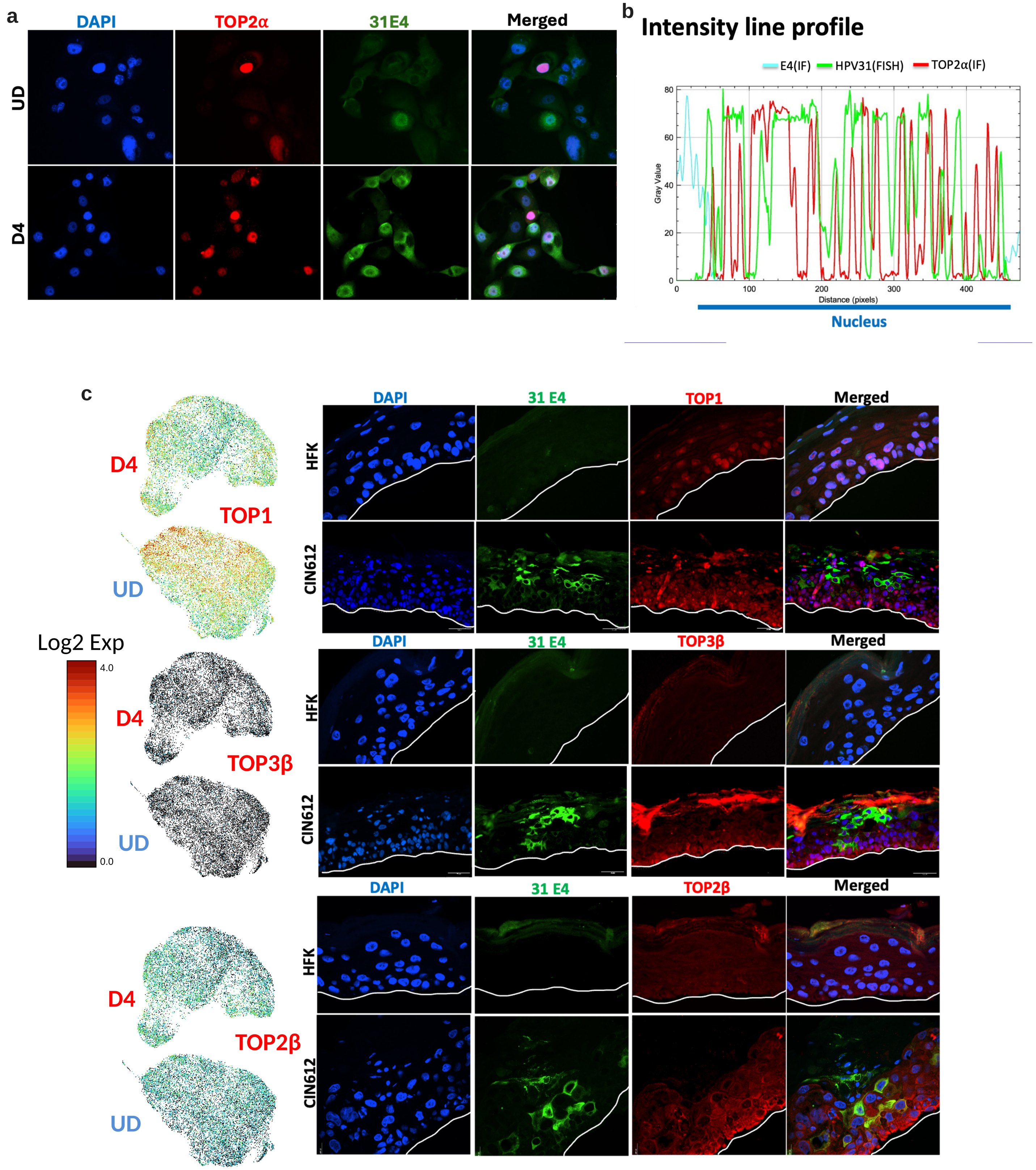


**Supplementary Figure 4 | TOP2α, but not other topoisomerases, is restricted to E1^E4-positive cells.** **a**, Co-immunofluorescence for DAPI, TOP2α and HPV31 E1^E4 (31E4) in undifferentiated (UD) and differentiated (D4) CIN612 monolayer cultures. **b**, Intensity line profile across a representative suprabasal nucleus showing E4 immunofluorescence, HPV31 FISH and TOP2α immunofluorescence signals. **c**, Co-immunofluorescence for HPV31 E1^E4 with TOP1, TOP3β or TOP2β in HFK and CIN612 organotypic rafts; the basement membrane is marked by a white line along with their respective UMAPs from single cell RNA sequencing (left).

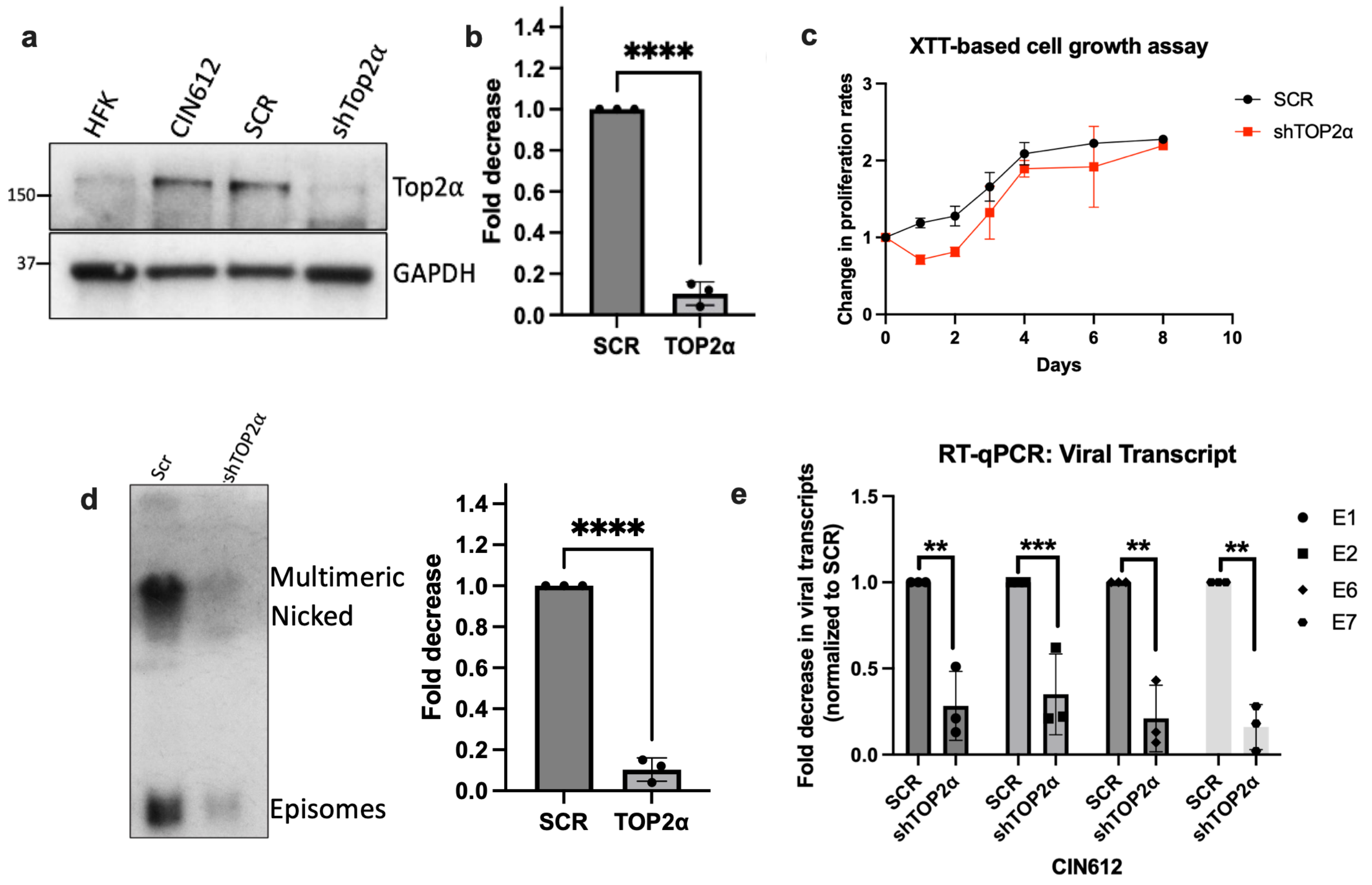


**Supplementary Figure 5 | Characterization of stable TOP2α knockdown CIN612 cells.** **a**, Immunoblot for TOP2α and GAPDH in HFK, CIN612, scrambled control (SCR) and stable shTOP2α CIN612 cells. **b**, Quantification of **a** (mean ± s.d., $n = 3$). **c**, XTT proliferation assay of SCR and stable shTOP2α CIN612 cells. **d**, Southern blot of HPV31 DNA in SCR and stable shTOP2α cells showing integrated, nicked and episomal species, with quantification of episomes (mean ± s.d., $n = 3$). **e**, RT–qPCR of *E1*, *E2*, *E6* and *E7* transcripts in SCR and stable shTOP2α CIN612 cells, normalized to GAPDH and to SCR. Two-sided unpaired Student *t*-test; $^{**}P \leq 0.01$, $^{***}P \leq 0.001$, $^{****}P \leq 0.0001$.

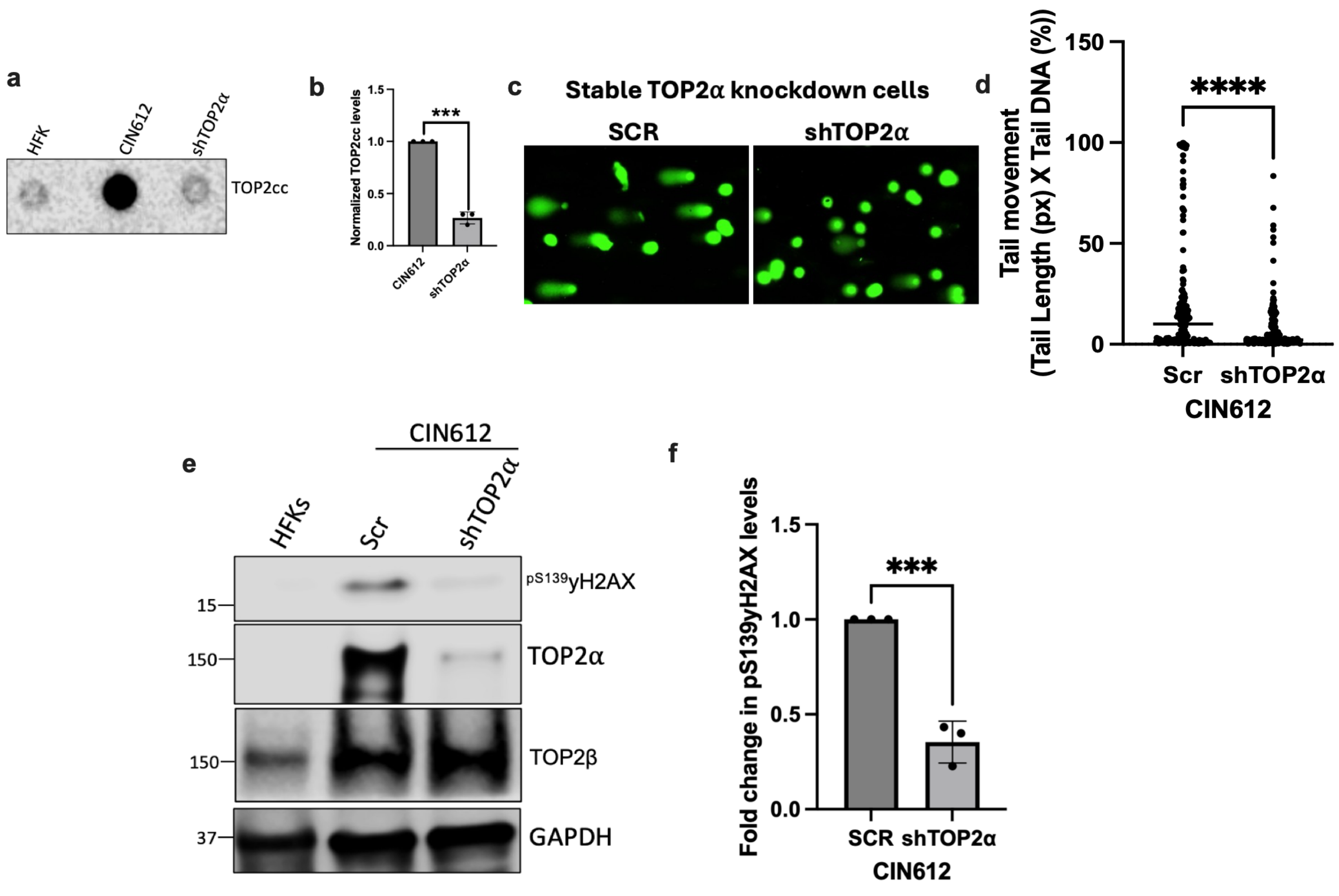


**Supplementary Figure 6 | Stable TOP2α knockdown reduces cleavage complexes, DNA breaks and γH2AX.** **a**, Slot blot of TOP2 cleavage complexes recovered by the RADAR assay from HFK, CIN612 and stable shTOP2α cells. **b**, Quantification of **a** (mean ± s.d., *n* = 3). **c**, Representative alkaline comet assay images from SCR and stable shTOP2α CIN612 cells. **d**, Tail moment for the conditions in **c**; at least 100 nuclei per condition. **e**, Immunoblot of HFK, SCR and stable shTOP2α CIN612 cells probed for pS139-γH2AX, TOP2α, TOP2β and GAPDH. **f**, Quantification of pS139-γH2AX from **e** (mean ± s.d., *n* = 3). Two-sided unpaired Student *t*-test (panels **b**, **f**) or two-sided Mann–Whitney *U* test (panel **d**); $***P \leq 0.001$, $****P \leq 0.0001$.
